# A human lung organoid co-culture model of early bone sarcoma metastasis reveals contact-dependent epithelial remodeling at the metastatic interface

**DOI:** 10.64898/2026.06.03.729838

**Authors:** Martha Magdalena Zylka, Tümay Capraz, Gaël Moquin-Beaudry, Christoph Hafemeister, Nikolina Bradaric, Lukas Watzke, Veveeyan Suresh, Ulrike Mann, Femke Ringnalda, Marco Wachtel, Matthias S. Dettmer, Corentin Thuillez, Matthias Farlik, Helena Sorger, Antonin Marchais, Karin Sanders, Marc van de Wetering, Markus G. Seidel, Bernadette Liegl-Atzwanger, Martin Metzelder, Polina Kameneva, Florian Halbritter, Heinrich Kovar, Branka Radic-Sarikas

## Abstract

Lung metastasis drives mortality across cancer types, yet how infiltrating tumor cells remodel the lung epithelium to establish metastatic niches remains poorly understood. Here we establish MESCUL (MEtastatic Sarcoma Co-CULture), a co-culture platform combining human lung organoids with patient-derived bone sarcoma cells to model early tumor–lung epithelial interactions in a physiologically relevant 3D system.

MESCUL reveals that direct tumor–epithelial contact induces rapid, reproducible lung epithelial remodeling across Ewing sarcoma (ES) and osteosarcoma (OS) models and multiple organoid donor backgrounds, which is contact-dependent and not recapitulated by paracrine signaling. Single-cell RNA sequencing identifies LIMES (Lung Interface Metastasis Signature), a shared transcriptional program encompassing focal adhesion assembly, matrix metalloprotease (MMP) upregulation, and emergence of a damage-associated transitional cell state, in both ES and OS. Mechanistically, tumor-derived fibronectin (FN1) engages epithelial integrin receptors to activate focal adhesion kinase (FAK), driving amphiregulin (AREG) induction and MMP-mediated remodeling; FN1 alone phenocopies this response, and FAK inhibition attenuates it, nominating the FN1–integrin–FAK–AREG axis as a candidate therapeutic vulnerability.

The LIMES program, identified through the MESCUL co-culture model, is spatially confined to the tumor–lung interface in patient metastases of both ES and OS, as demonstrated by spatial transcriptomics across nine patients. Masson’s trichrome staining of matched patient sections reveals pronounced collagen deposition in the peri-tumoral lung parenchyma, consistent with LIMES acting upstream of a wound-healing cascade that proceeds to structural fibrotic remodeling in patient tissue.

Together, these findings establish the lung epithelium as an active participant in metastatic colonization, characterize a pharmacologically targetable, spatially restricted epithelial remodeling response at the bone sarcoma–lung interface across OS and ES, and introduce MESCUL as a tractable 3D platform for investigating lung metastasis, with possible implications for tumor types beyond bone sarcomas.

## Background

Metastasis is the principal driver of cancer-related mortality, responsible for the vast majority of cancer deaths worldwide [1]. The lung is one of the most frequent sites of distant metastasis across tumor types [2], and pulmonary colonization represents a critical and often fatal step in disease progression. This is particularly true for pediatric bone sarcomas patients of whom about 20 to 25% present with metastasis at diagnosis [3,4]. Ewing sarcoma (ES) and osteosarcoma (OS) are highly aggressive malignancies, with five-year overall survival rates of 70–80% for patients with localized disease but only approximately 30% for those presenting with metastases [5,6]. In both entities, the lung is the dominant metastatic site [7,8] and pulmonary failure the principal cause of treatment-related mortality. Despite advances in systemic therapy, survival for patients with metastatic bone sarcoma has not improved meaningfully for several decades [9], underscoring the urgent need for a deeper mechanistic understanding of pulmonary metastatic homing and colonization.

Despite the clinical urgency, the molecular events during metastatic homing to the lung remain incompletely understood. In OS, a wound-healing and fibrotic transcriptional program has been identified at the tumor–lung interface in murine models [10]; whether this response translates to human disease, and whether an analogous program operates in ES, remain unknown. A fibrosis-like or epithelial remodeling program at the ES–lung interface has not previously been described.

Mechanistic exploration of early tumor–epithelium interactions has been hampered by the limitations of available experimental systems. Conventional in vitro models based on established cell lines lack the cellular heterogeneity and three-dimensional architecture of the human lung epithelium, while murine models do not fully recapitulate human-specific epithelial biology [11,12].

Here, we establish MESCUL (MEtastatic Sarcoma Co-CULture) - a human lung organoid-based co-culture platform for mechanistic interrogation of bone sarcoma - lung epithelial interactions using patient material. Lung organoids are self-organizing three-dimensional epithelial cultures that recapitulate the complexity of the airway epithelium [13]. Using MESCUL, we characterized a previously undescribed epithelial transcriptional program, the Lung Interface MEtastasis Signature (LIMES), activated upon direct bone sarcoma-epithelium contact, promoting active lung parenchymal remodeling. Using spatial transcriptomics, we confirm our ex vivo findings from MESCUL at the tumor–lung interface in bone sarcoma patient metastases. Together, our findings establish the lung epithelium as an active partner in crime in metastatic colonization in bone sarcomas, and position MESCUL as a versatile platform for mechanistic and therapeutic investigation of lung homing.

## Results

### Patient-derived pediatric lung organoids recapitulate epithelial diversity and enable in vitro modeling of early metastatic homing

To establish a physiologically relevant platform for studying early metastatic tumor–epithelium interactions, we generated lung organoids from non-malignant pediatric lung tissue resected adjacent to either benign malformations or metastatic lesions, with organoids from all six donors displaying comparable morphologies, confirming robustness of the establishment protocol (Figure 1a, Supplementary Figure 1a, Supplementary Table 1). Immunostaining confirmed the presence of basal, club, ciliated, and goblet cell populations, and differentiated organoids maintained the key physiological function of mucociliary activity (Figure 1b, Supplementary Movie 1). To resolve the epithelial composition at single-cell resolution, we performed single-cell RNA-sequencing (scRNA-seq) on organoids from three donors (L3, L-EWS-050, L-OS-060) and projected them into a joint Uniform Manifold Approximation and Projection (UMAP) (Figure 1c). Manual cell type annotation based on a curated set of in vitro-adapted marker genes revealed the presence of alveolar type 2 (AT2), AT2 progenitor, pre-alveolar type-1 transitional (PATS), and alveolar type 1 (AT1) cells, alongside the expected airway populations, including basal, secretory, and multiciliated cells (Figure 1d). PATS represent a transcriptionally distinct transitional state bridging the AT2-to-AT1 differentiation trajectory and are associated with alveolar injury response [14–17]. Notably, the model captured AT1 cells, which are rarely maintained in organoid culture [18], indicating preservation of distal lung identity and active alveolar lineage progression beyond what is typically achieved in airway-focused systems. Organoid-derived cells mapped broadly across epithelial populations of the Human Lung Cell Atlas (LungMAP, [19]), with overlap across both airway and alveolar lineages, further confirming that the model faithfully captures the cellular heterogeneity of the human lung epithelium (Supplementary Figure 1b, c). We therefore used pediatric lung organoids to study the response of distinct lung epithelial cell types to tumor metastatic cues ex vivo.

**Figure 1.**
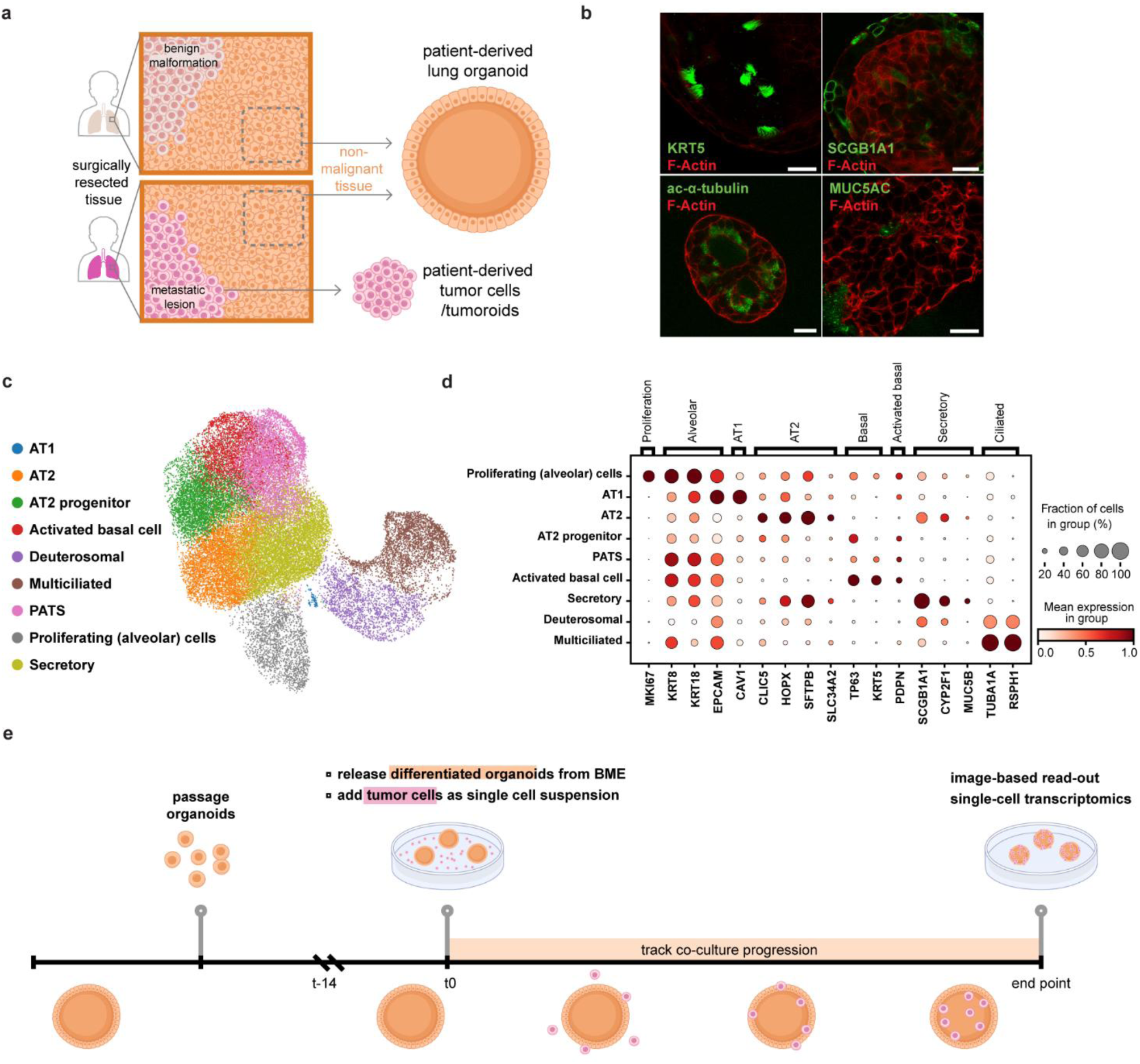
Patient-derived pediatric lung organoids recapitulate epithelial diversity and enable in vitro modeling of early metastatic homing. **a,** Schematic of non-malignant pediatric lung tissue sources used for organoid generation and, when applicable, tumor cell isolation, depicting sampling regions adjacent to benign malformations or metastatic lesions. **b**, Immunostaining of differentiated lung organoids confirming the presence and spatial organization of major epithelial cell types: basal cells (KRT5), club cells (SCGB1A1), ciliated cells (acetylated-α-tubulin), and goblet cells (MUC5AC). Cell-type markers are shown in green and F-actin in red. Scale bar, 20 µm. **c**, UMAP embedding of scRNA-seq data from pediatric lung organoids across multiple donors, with cell type annotations based on a curated set of in vitro-adapted marker genes. **d**, Dot plot depicting the expression of canonical marker genes across manually annotated epithelial cell populations, illustrating the transcriptional basis for cell type assignments shown in c. Dot size indicates the fraction of cells in the cluster expressing the gene; color intensity indicates mean expression level. **e**, Schematic of the MESCUL co-culture approach. Differentiated organoids were released from BME and combined with tumor cells as a single-cell suspension. Co-cultures were maintained for up to three weeks with regular imaging until endpoint transcriptomic and molecular analysis.

To model early tumor–lung epithelial interactions, we developed MESCUL (Metastatic Sarcoma Co-Culture), a co-culture system in which lung organoids were first expanded and differentiated embedded in basement membrane extract (BME), then released and combined with ES or OS cells as a single-cell suspension (Figure 1e). Tumor cells derived from either freshly isolated primary patient material or patient-derived tumoroids were labeled for live tracking using stable GFP expression or CellTrace™, enabling reliable discrimination from organoid-derived cells throughout the assay (Supplementary Table 1). MESCUL co-cultures were maintained for up to three weeks and imaged at regular intervals, and samples were collected at the endpoint to enable transcriptomic and molecular characterization of the evolving tumor-epithelium interactions.

### Live-cell imaging reveals rapid Ewing sarcoma cell infiltration and initiation of lung epithelial remodeling

To directly visualize tumor–epithelium interactions in real time, we monitored MESCUL co-cultures of GFP-labeled patient-derived ES cells (ES-010) with lung organoids by live-cell imaging. Early after co-culture initiation, tumor cells established contact with the outer organoid surface and began to enter and intercalate within the lung epithelium (Figure 2a, Supplementary Movie 2). Active infiltration was confirmed by maximum-intensity projections of the GFP channel, which revealed a transient reduction in fluorescence intensity as tumor cells traversed the outer epithelial layer, followed by signal recovery upon successful entry (Figure 2b). Strikingly, infiltration was accompanied by marked organoid remodeling, including loss of 3D architecture, adhesion to the culture plate surface, and pronounced epithelial spreading into a flattened morphology reminiscent of epithelial-to-mesenchymal transition (EMT), none of which occurred in matched mono-cultures (Figure 2c). Confocal imaging of an independent co-culture replicate corroborated these findings, revealing tumor cells intercalating between lung epithelial cells and disrupting organoid architecture, with epithelial cells acquiring mesenchymal properties throughout (Figure 2f, g). Quantification of organoid area from live-cell imaging, which enabled tracking of the same organoids over time, confirmed a robust and significant increase in co-cultured organoids (p=0.0312, Wilcoxon matched-pairs signed-rank test, n=6), whereas mono-cultured organoids showed no statistically significant change over the same interval (n=3) (Figure 2d, e), establishing organoid area as a quantifiable readout of tumor-induced epithelial remodeling.

**Figure 2.**
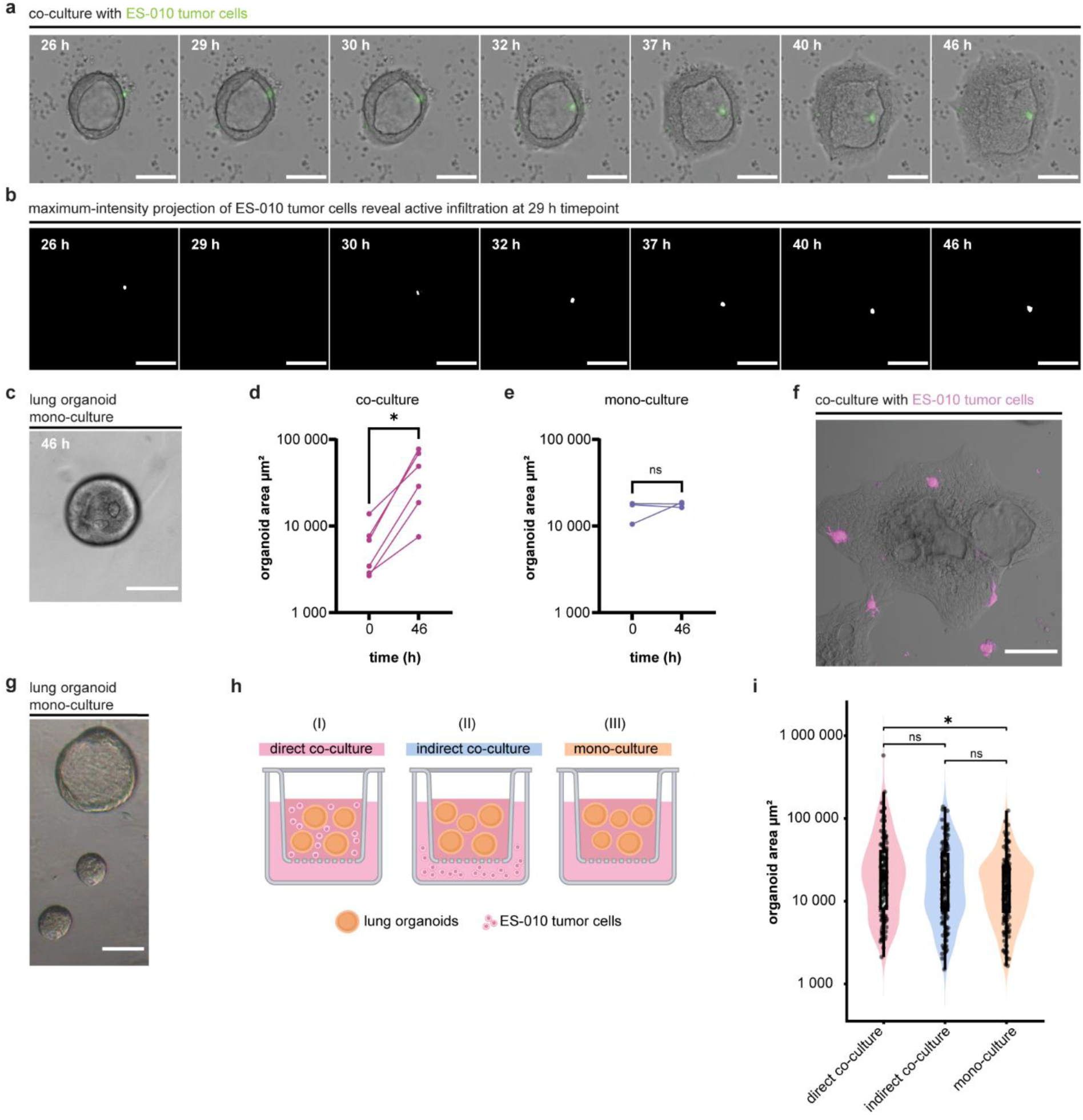
Ewing sarcoma cells actively infiltrate lung organoids and induce epithelial remodeling dependent on direct tumor–epithelial contact. **a,** Representative brightfield time-lapse series of MESCUL co-culture of GFP-labeled ES-010 cells, showing progressive tumor cell infiltration and organoid remodeling over 46 hours. Scale bar, 100 µm. **b,** Maximum-intensity projections of the GFP channel from the same time-lapse series shown in a, illustrating transient fluorescence attenuation as tumor cells traverse the outer epithelial layer. Scale bar, 100 µm. **c,** Representative brightfield image of a lung organoid mono-culture at 46 hours, shown as a reference for unperturbed organoid morphology. Scale bar, 100 µm. **d, e,** Organoid area (µm²) of individual lung organoids tracked from t=0 to t=46h in co-culture (d) and mono-culture (e) conditions, including the representative organoid shown in panel a. Each line represents one organoid tracked within a single well across both timepoints (one experiment; co-culture n=6, mono-culture n=3). Statistical comparisons between t=0 and t=46h were performed using the Wilcoxon matched-pairs signed-rank test (two-tailed). *p<0.05, ns = not significant. **f,** Representative confocal image of a lung organoid co-culture with ES-010 tumor cells (magenta), showing tumor cell intercalation and disruption of organoid architecture. Scale bar, 100 µm. **g,** Representative confocal image of lung organoid mono-culture. Scale bar, 100 µm. **h,** Schematic of the three experimental conditions used to dissect the mode of tumor–epithelium interaction: (i) direct co-culture, (ii) indirect co-culture via a permeable membrane, and (iii) mono-culture. **i,** Organoid area (µm²) of lung organoids cultured under direct co-culture, indirect co-culture, and mono- culture conditions. Each dot represents an individual organoid measurement (direct: n=134, indirect: n=144, mono-culture: n=154; 3 independent biological replicates, 2 technical replicates per condition, 15–40 organoids per well). Individual organoid measurements are nested within wells and biological replicates; statistical comparisons were performed using a linear mixed effects model with Tukey-corrected pairwise comparisons. *p<0.05, ns = not significant.

Together, these findings demonstrate that patient-derived ES cells actively infiltrate lung organoids, inducing quantifiable structural remodeling and an EMT-like response in the lung epithelium.

### Lung organoid epithelial remodeling is contact-dependent and cannot be triggered by paracrine signaling alone

To determine whether the epithelial remodeling phenotype is mediated by tumor-derived soluble factors or requires direct physical interaction, we compared three conditions: (i) direct co-culture, permitting unrestricted tumor–epithelium contact; (ii) indirect co-culture, in which organoids and tumor cells shared medium across a permeable membrane but remained physically separated; and (iii) mono-culture controls (Figure 2h). Lung organoids exposed to direct co-culture displayed a significant increase in organoid area compared to mono-culture (p=0.0111) (Figure 2i). In contrast, organoid size in the indirect condition did not differ significantly from mono-culture (p=0.5083), demonstrating that soluble factors alone do not suffice to trigger epithelial remodeling and that direct tumor–epithelial contact is required for this response.

### Lung epithelial remodeling is reproducibly induced across Ewing sarcoma and osteosarcoma models and multiple organoid donor backgrounds

To determine whether the observed remodeling phenotype extends beyond ES-010, we systematically tested multiple additional ES and OS models across lung organoids derived from three independent donor backgrounds, including tissue resected adjacent to benign lung malformation, OS lung metastasis, and ES lung metastasis. Large-scale quantification across four co-culture experiments, encompassing 2,812 individually segmented organoids, revealed an increase in organoid area relative to mono-culture controls across all tested organoid–tumor cell combinations (Figure 3a–d). In organoids derived from benign lung tissue, co-culture with ES-046 significantly increased organoid area (∼1.9-fold), while ES-010 showed a directionally consistent trend (p = 0.071, ∼1.6-fold; Figure 3a). This pattern was reproduced in organoids derived from OS lung metastasis, where both ES-010 (∼1.2-fold) and ES-046 (∼1.5-fold) co-culture significantly increased organoid area (Figure 3b). Co-culture with ES-016 in organoids derived from ES lung metastasis confirmed organoid tissue remodeling phenotype across a distinct patient background (∼5.1-fold; Figure 3c). Extending these observations to OS, OS-080 co-culture produced a comparable increase in organoid area in benign-derived L3 organoids (∼2.0-fold; Figure 3d). Live-cell imaging captured the dynamic progression of this response, showing rapid tumor cell contact with lung organoids followed by progressive disruption of the organoid epithelium and attachment to the culture surface, as observed with EWS-040, EWS-050, and OS-060 tumor cells (Figure 3e–g). Taken together, these results demonstrate that sarcoma-induced epithelial remodeling is a reproducible and conserved response across the tumor specimen and organoid donor backgrounds tested, observed across both ES and OS.

**Figure 3.**
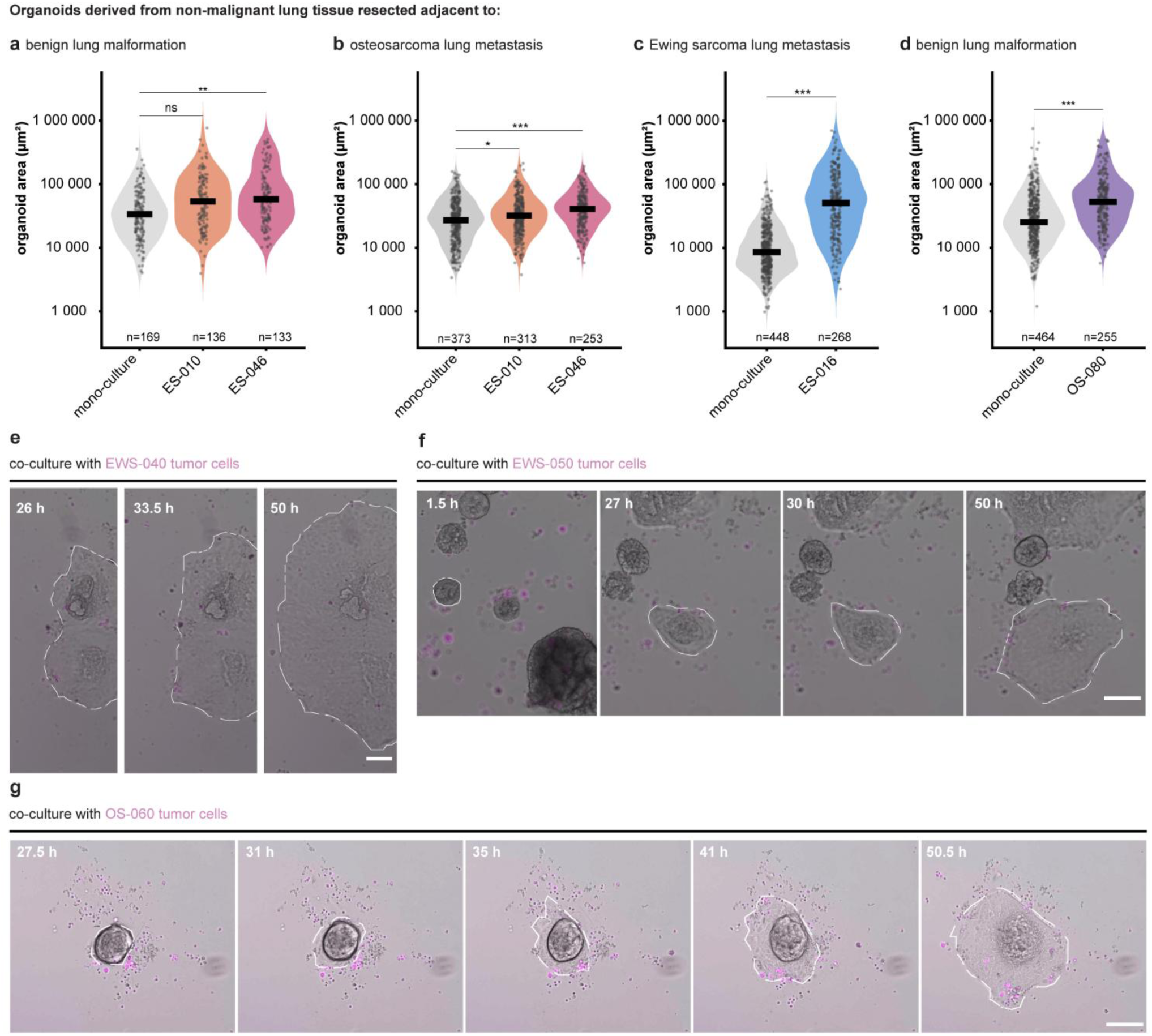
Sarcoma-induced lung epithelial remodeling is reproducible across multiple tumor specimen and organoid donor backgrounds. **a–d**, Violin plots showing organoid area (µm²) of patient-derived lung organoids in mono-culture or co-culture with the indicated tumor cells. Organoids were derived from non-malignant lung tissue resected adjacent to benign lung malformation (L3; a, d), OS lung metastasis (L-OS-060; b), or ES lung metastasis (L-EWS-050; c). **a, b,** Tumor specimen: ES-010 and ES-046 (matched primary and relapsed tumor from the same patient; **c,** ES-016 (independent patient; **d,** OS-080. Each data point represents one individually segmented organoid; n per condition is indicated below each violin. Horizontal bars indicate the median. Each experiment comprised 4 technical replicate wells per condition and represents one biological replicate. Organoid area was log10-transformed prior to statistical analysis and compared using a linear mixed-effects model (log10(area) ∼ condition + (1 | well)), with pairwise contrasts corrected by Tukey’s method. Effect sizes: a ES-010: ∼1.6-fold increase, p = 0.071; ES-046: ∼1.9-fold increase, p = 0.009. b ES-010: ∼1.2-fold increase, p = 0.038; ES-046: ∼1.5-fold increase, p < 0.001. c ES-016: ∼5.1-fold increase, p < 0.001. d OS-080: ∼2.0-fold increase, p < 0.001. All fold-changes refer to median organoid area in co-culture relative to mono-culture. * p < 0.05, ** p < 0.01, *** p < 0.001; ns, not significant. **e–g,** Representative brightfield time-lapse series of MESCUL co-cultures of **e,** EWS-040, **f,** EWS-050 and **g,** OS-060 tumor cells (magenta), showing progressive tumor cell contact, organoid flattening, and epithelial spreading over time. Scale bars, 100 µm.

### Bone sarcoma contact induces a conserved maladaptive transcriptional response in the lung epithelium

Having established that lung epithelial remodeling is a conserved and contact-dependent response to bone sarcoma infiltration, we next sought to characterize its transcriptional basis. We performed scRNA-seq (10x Genomics 3′, v 3.1) on lung organoids co-cultured with three ES (ES-010, ES-016, ES-080) or two OS tumor samples (OS-060, OS-080) and compared them to matched mono-culture controls across the five independent co-culture experiments (Figure 4a). To identify the core epithelial response, we restricted analysis to genes differentially expressed in the same direction across all five conditions, yielding a tumor-entity- and sample-independent convergent transcriptional program (DESeq2 Wald-test on pseudobulks)(Supplementary Table 2). From these conserved changes, we derived a curated gene signature encompassing extra-cellular matrix (ECM) remodeling, focal adhesion, and cytoskeletal programs, which we termed the *Lung Interface Metastasis Signature* (LIMES). The LIMES genes - including the ECM protease *MMP1*, the actin-bundling protein *FSCN1*, the stress keratin *KRT6A*, and the EGFR (epidermal growth factor receptor) ligands *AREG* (amphiregulin) and *EREG* (epiregulin) - were uniformly upregulated across all co-culture conditions, indicating a robust epithelial transcriptional response to bone sarcoma contact irrespective of tumor entity or sample origin (Figure 4a; Supplementary Table 3).

**Figure 4.**
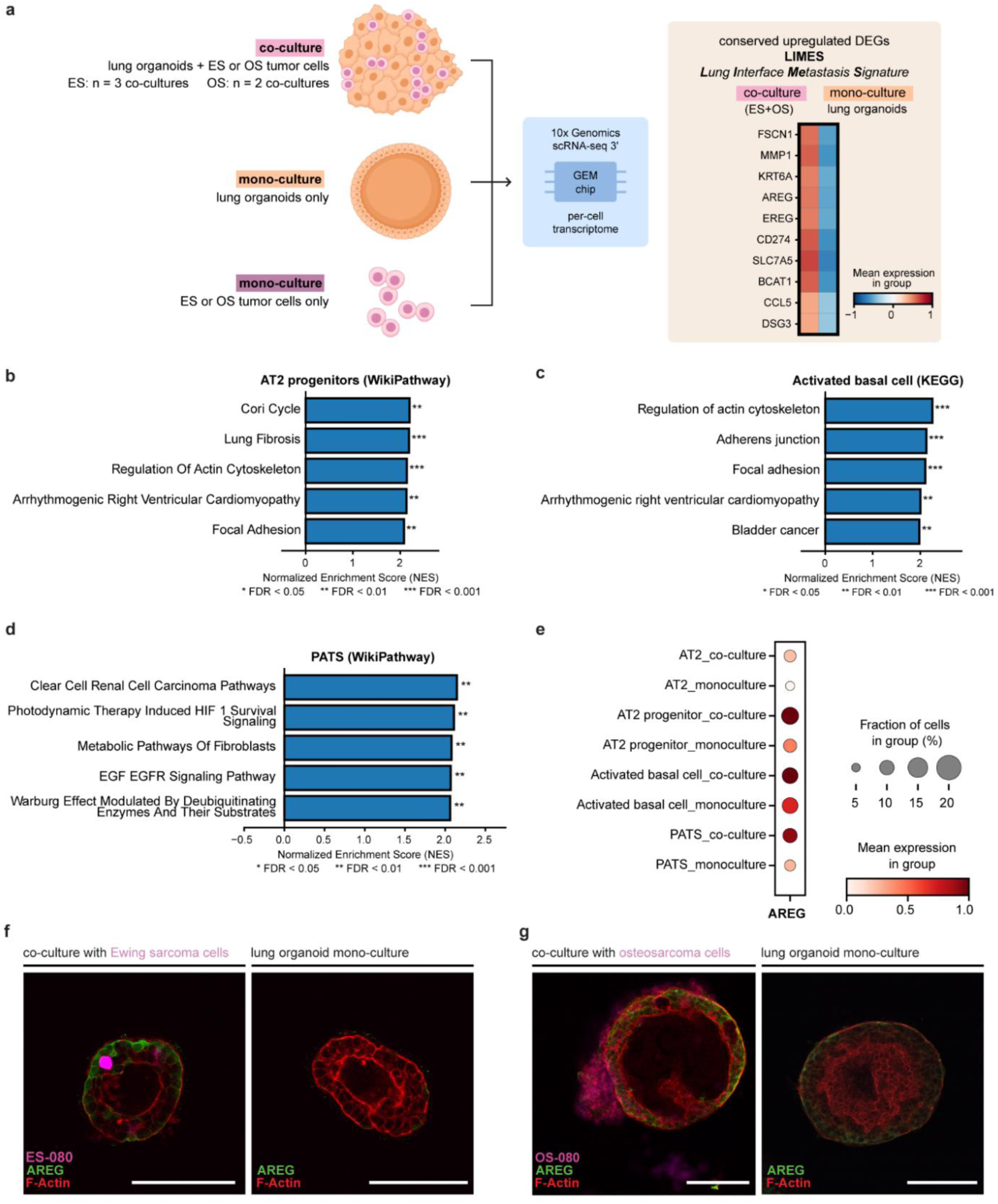
Sarcoma contact induces a conserved maladaptive transcriptional response in the lung epithelium. **a,** Experimental design for scRNA-seq. Lung organoids were co-cultured with ES (n = 3 co-cultures) or OS (n = 2 co-cultures) tumor cells, or maintained as mono-cultures, and processed for 10x Genomics scRNA-seq. The heatmap shows the conserved upregulated differentially expressed genes shared between ES and OS co-cultures relative to mono-culture controls, constituting the Lung Interface Metastasis Signature (LIMES). **b–d,** GSEA of co-culture-upregulated genes in AT2 progenitor cells using WikiPathway annotations (b), in activated basal cells using KEGG annotations (c), and in PATS using WikiPathway annotations (d). Bar length indicates normalized enrichment score (NES); * indicate significance levels (* FDR < 0.05, ** FDR < 0.01, *** FDR < 0.001 **e,** Dot plot showing AREG expression across epithelial cell types in co-culture and mono-culture conditions. Dot size indicates the fraction of cells expressing AREG; color intensity indicates mean expression level. **f, g,** Confocal images of lung organoids co-cultured with ES (f) or OS (g) tumor cells, immunostained for AREG (green) and F-actin (red). Tumor cells are shown in magenta. Scale bars, 100 µm.

Cell-type-resolved Gene set enrichment analysis (GSEA) further revealed cell-type-specific transcriptional programs reflecting distinct injury states (Supplementary Table 4). AT2 progenitor cells showed enrichment of a lung fibrosis gene set alongside focal adhesion pathway terms (Figure 4b). Activated basal cells exhibited the strongest enrichment of focal adhesion assembly and adherens junction programs (Figure 4c), consistent with the observed mechanosensitive, contact-activated state. PATS showed induction of fibroblast metabolic pathways, HIF-1 survival signaling, and EGF–EGFR signaling (Figure 4d), consistent with an injury-arrested transitional state exhibiting acquired fibroblast-like transcriptional features, recently described to be the onset of fibrotic disease [14–17].

Among the LIMES genes, the EGFR ligand *AREG* emerged as a potential marker of the co-culture state: its upregulation was concentrated in transitional epithelial populations - AT2 progenitors, activated basal cells, and PATS - with negligible expression in mono-culture controls across all five experiments (Figure 4e). Given AREG’s established roles in epithelial repair, lung fibrosis, and tumor progression [20–22], its selective induction in injury-related cell states further corroborates the pro-fibrotic, maladaptive character of the epithelial response to tumor cell contact. Immunofluorescence staining confirmed AREG upregulation in lung organoids co-cultured with either ES and OS cells relative to matched monoculture controls, with signal distributed broadly across the organoid epithelium and largely reduced in tumor cells (Figures 4f,g, Supplementary Figure 2a,b) validating AREG induction as a shared epithelial response to sarcoma cell contact at the protein level across both entities.

### Tumor-derived FN1 activates a contact-dependent integrin–FAK axis that drives proteolytic remodeling of the lung epithelium

To identify the signals driving the LIMES program, we examined the transcriptional landscape of co-cultured tumor cells. Among upregulated genes in co-cultured ES-010 cells, *fibronectin-1* (*FN1*) was identified as a tumor-derived candidate, showing significant transcriptional upregulation relative to monoculture controls. *FN1* upregulation was accompanied by coordinated induction of *ITGAV* and *ITGB1*, encoding the integrin subunits αv and β1 that heterodimerize to form the αvβ1 complex - a canonical FN1-binding integrin competent for cell adhesion and extracellular matrix engagement. (Figure 5a) [23,24]. Immunofluorescence staining confirmed FN1 protein deposition specifically at the tumor–organoid interface in both ES (ES-010) and OS (OS-080) co-cultures, with minimal scattered signal in matched tumor cell and lung organoid monocultures (Figures 5b, c).

**Figure 5.**
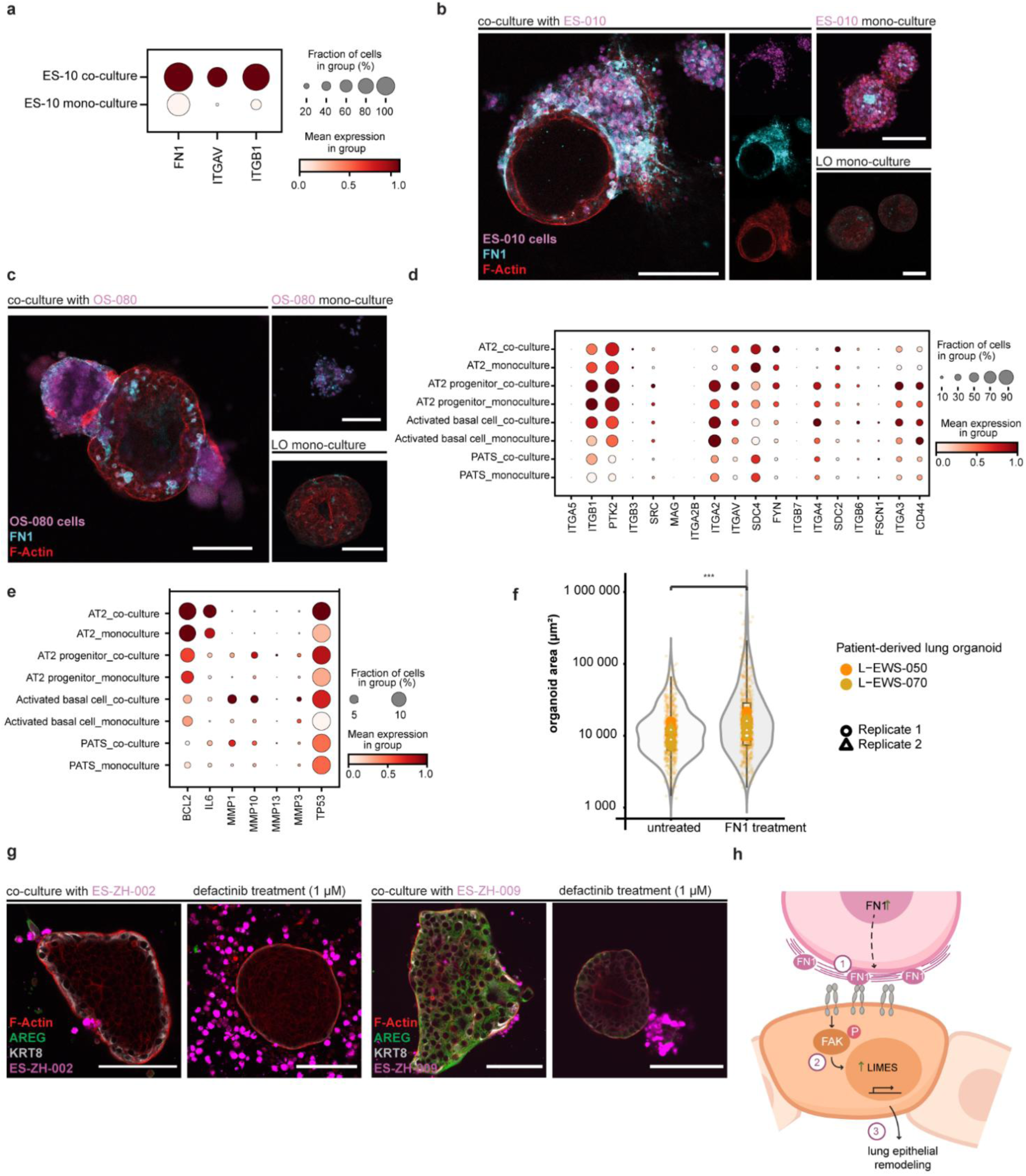
Tumor-derived FN1 engages a contact-dependent integrin–FAK axis that co-opts the lung epithelium into proteolytic self-remodeling. **a,** Dot plot showing expression of FN1, ITGAV, and ITGB1 in ES-010 cells recovered from MESCUL co-culture versus ES-010 monoculture. **b,** Immunofluorescence images of MESCUL co-culture with ES-010 cells (left; large panel with split-channel insets showing tumor cells, FN1, and F-Actin separately) and matched ES-010 tumor cell monoculture and lung organoid monoculture controls (right). Staining: ES-010 cells (magenta), FN1 (cyan), F-Actin (red). Scale bars, 100 µm. **c,** Immunofluorescence images of lung organoids co-cultured with OS-080 OS cells (left) and matched OS-080 tumor cell monoculture and lung organoid monoculture controls (right). Staining: OS-080 cells (magenta), FN1 (cyan), F-Actin (red). Scale bars, 100 µm. **d,** Dot plot showing expression of FN1 receptor candidates across annotated lung epithelial cell populations in co-culture versus monoculture conditions. Dot size represents the fraction of cells expressing the gene; color intensity represents scaled mean expression (scale 0.0–1.0). **e,** Dot plot showing expression of NicheNet-predicted FN1 downstream target genes across annotated lung epithelial cell populations in co-culture versus monoculture conditions. Dot size represents the fraction of cells expressing the gene; color intensity represents scaled mean expression (scale 0.0–1.0). **f,** Violin plots showing organoid area (µm²) of patient-derived lung organoids treated with fibronectin (FN1, 64 µg/mL) or left untreated for 6 days post-seeding. Two independent patient-derived lung organoid lines (L-EWS-050, L-EWS-070) were assessed across two independent experiments (4 wells per condition in replicate 1; 3 wells per condition in replicate 2). Individual data points represent single organoid area measurements, colored by patient line; open symbols indicate per-well means (circles: replicate 1; triangles: replicate 2). Organoid area was measured log₁₀-transformed prior to statistical analysis. To account for the non-independence of organoids nested within wells, and wells nested within patient-by-experiment batches, a linear mixed effects model was fitted of the form log₁₀(area) ∼ condition + (1 | patient) + (1 | batch/well), where batch denotes the patient-by-experiment combination (lme4; Kenward-Roger degrees of freedom, lmerTest). *** p < 0.001 (FN1 vs. untreated). **g,** Immunofluorescence images of lung organoids co-cultured with ES-ZH-002 (left) or ES-ZH-009 (right) ES cells under control conditions or following treatment with 1 µM defactinib, stained for F-Actin (red), AREG (green), KRT8 (white), and tumor cells (magenta). n=2 independent co-cultures across two patient-derived ES lines. Scale bars, 100 µm. **h,** Schematic illustration of the proposed FN1–integrin–FAK signaling axis at the sarcoma–lung epithelial interface. (1) Tumor-derived FN1 is deposited at the contact interface and engages integrin receptors on lung epithelial cells. (2) Integrin engagement activates FAK, driving downstream upregulation of LIMES program (3) Collectively, these signals drive lung epithelial remodeling at the metastatic niche.

On the epithelial side, using NicheNet’s database of ligand-receptor interactions, we identified the FN1 targeted receptors *ITGB1, ITGAV*, *ITGA3*, *PTK2*, *CD44*, and *SDC4*, with expression of these receptors confirmed across co-cultured lung epithelial populations - most prominently in AT2 progenitor and activated basal cell populations (Figure 5d). *ITGB1* and *ITGAV* were among the identified receptors, mirroring their co-culture-induced upregulation in ES-010 tumor cells. *ITGA3* has been independently identified as a principal epithelial FN1 receptor at the OS–lung interface [25], and *PTK2* enrichment in AT2 progenitor co-cultures is consistent with its role as a downstream transducer of integrin engagement [26]. Together, this receptor profile positions the lung epithelium to receive and transduce the tumor-derived FN1 signal at the point of direct contact. In addition, using NicheNet’s database we identified the stromelytic metalloproteases *MMP1*, *MMP3*, and *MMP10*, alongside *BCL2* and *IL6* as downstream targets of FN1 signaling - a proteolytic and pro-inflammatory effector profile consistent with the observed active extracellular matrix remodeling at the lung organoid metastatic niche (Figure 5e).

To functionally validate the sufficiency of tumor-derived FN1 to drive epithelial remodeling, lung organoid monocultures were treated with human-plasma derived fibronectin, which induced a significant increase in organoid area across two independent patient-derived organoid lines, demonstrating that FN1 alone is sufficient to induce organoid remodeling independently of other potential tumor-derived signals (Figure 5f, Supplementary Figure 2c,d). Given that FN1–integrin engagement canonically activates FAK through integrin clustering and β1-tail phosphorylation [24,27], and consistent with NicheNet predicting *PTK2*, the gene encoding the focal adhesion kinase FAK, as part of the FN1-driven epithelial response, we asked whether FAK inhibition would attenuate the remodeling phenotype. Treatment with the selective FAK inhibitor defactinib (1 µM; dose selected based on organoid and tumor cell viability assays, Supplementary Figure 2e,f) reduced organoid area in direct co-culture compared to vehicle control and markedly reduced AREG expression - as confirmed at the protein level (Figure 4f,g) - alongside depletion of KRT8⁺ transitional cells, a damage-associated epithelial state characteristic of the PATS population that accumulates during aberrant alveolar repair [14,15], across two independent ES co-cultures (Figure 5g) - demonstrating that FAK activity is required for contact-dependent epithelial remodeling and AREG induction downstream of integrin engagement. To identify candidate upstream regulators of tumor FN1 expression, we profiled the secretome of lung organoid monocultures, reasoning that lung epithelial-conditioned medium is sufficient to induce tumor FN1 deposition in osteosarcoma cells [10]. GDF-15 (Growth-differentiation-factor 15) emerged as the most abundantly secreted factor, suggesting it as a candidate epithelial priming signal (Supplementary Figure 2g). GDF-15 is a divergent TGF-β superfamily member previously shown to be upregulated in osteosarcoma pulmonary metastases and to promote osteosarcoma invasion through activation of TGF-β/SMAD2/3 signaling [28]. As TGF-β/SMAD2/3 signaling is a well-established transcriptional inducer of FN1 [29], these data collectively nominate lung epithelial GDF-15 as a candidate upstream regulator that may prime arriving tumor cells toward an FN1-expressing state prior to direct epithelial contact, establishing the ligand supply required for subsequent contact-dependent integrin engagement. The proposed mechanistic sequence - tumor-derived FN1 engaging epithelial integrins at the contact interface, and FAK-dependent downstream activation of MMP1/2 and AREG driving lung epithelial remodeling - is summarized in Figure 5h.

### The LIMES program is spatially confined to the tumor–lung interface and accompanied by fibrotic tissue remodeling in Ewing sarcoma and osteosarcoma lung metastases

To assess whether the LIMES program identified in our patient-derived metastasis model is recapitulated in patient lung metastases, we performed spatial transcriptomics (10x Genomics Visium) on matched primary tumors and lung metastases from nine patients (three ES, six OS)(Supplementary Table 5). For two ES patients (ES2, ES3), multiple spatially independent metastatic lesions from distinct lung locations were available, yielding 12 metastatic sections in total (Supplementary Figure 3a,b). Spatial mapping of the LIMES gene signature revealed enrichment at the immediate tumor–lung interface in metastatic lesions of both entities, whereas signal in corresponding primary tumors was weaker and lacked spatial coherence relative to the tumor boundary (Figures 6a, b, Supplementary Figure 4). To quantify this distribution, we measured the distance to the tumor front, expressed as the minimal number of Visium spot “hops”, at which LIMES signal was the highest on average, herein named peak signal hop, across all 12 metastatic sections. This analysis confirmed a stromal-side, boundary-proximal peak in 10 of 12 individual metastatic sections (Figure 6c), whereas matched primary tumor sections showed no comparable enrichment pattern, with peak signal distributed without consistent spatial organization relative to the tumor boundary (Figure 6d). In a distance-binned expression plot across all samples, a clear trend of signal increase at or immediately beyond the tumor boundary was observed for the lung metastasis patient cohort, indicating that LIMES induction is spatially concentrated at the tumor-lung interface (Figure 6e,f). Together, these findings confirm that the epithelial remodeling program identified in our co-culture system is also observed at the tumor-lung interface in patient metastases, providing in vivo transcriptomic validation of the LIMES signature.

**Figure 6.**
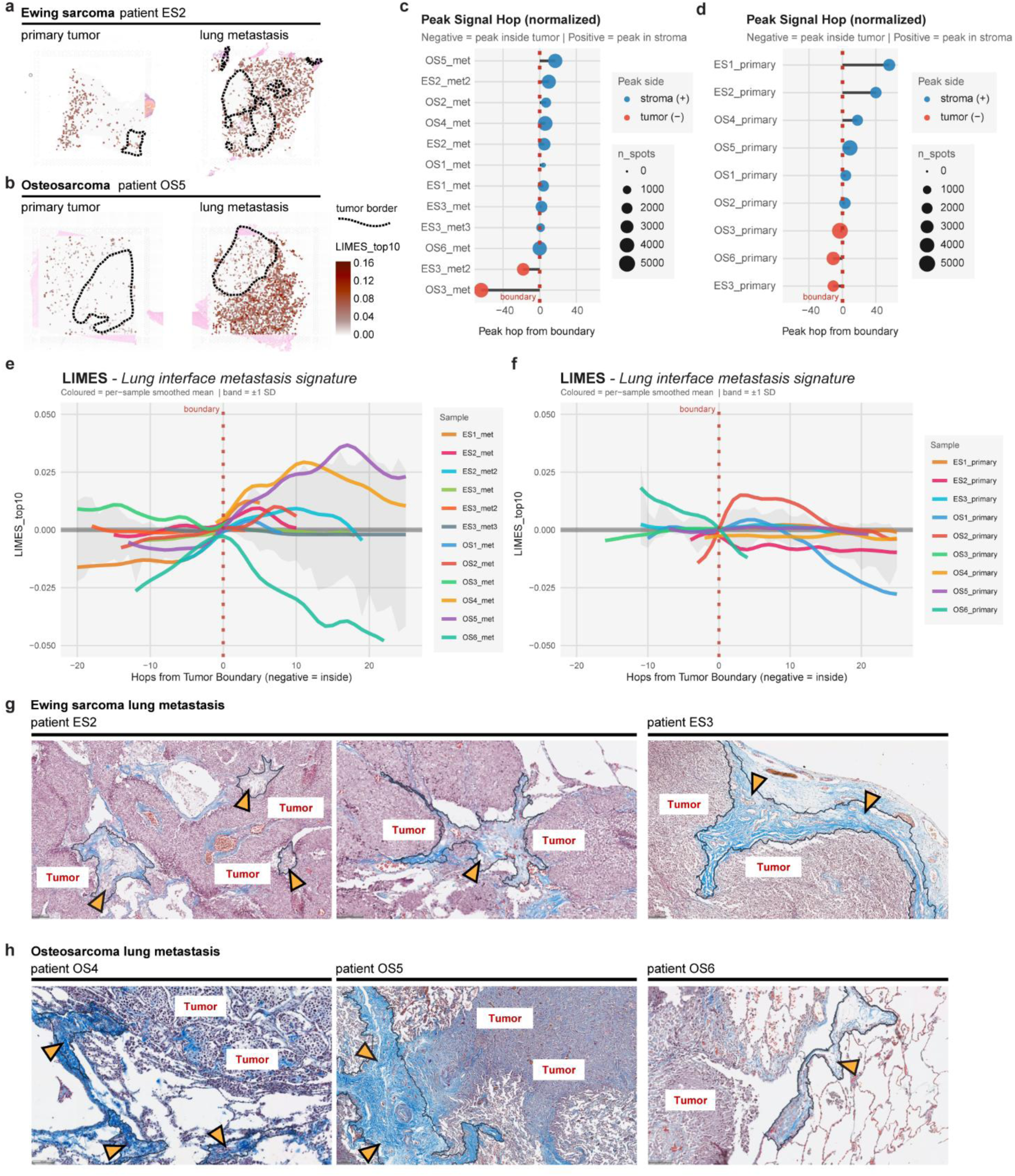
Spatial validation of the LIMES program and downstream effects at the tumor–lung interface in patient metastases. **a,** 10x Visium spatial transcriptomics section from ES patient ES2 (primary tumor and matched lung metastasis). LIMES (top10) UCell score is plotted per spot; tumor boundary is annotated with a dashed line. Color scale indicates normalized UCell score.**b,** As in a, for OS patient OS5 (primary tumor and matched lung metastasis).**c,** Peak signal hop analysis across 12 lung metastatic sections. Distance is expressed as the minimal number of spot hops to a tumor boundary spot on the hexagonal geometry of the Visium array, with center-to-center distance between spots of 100 µm. Dot position on the x-axis indicates the hop of maximum boundary-normalized LIMES score per section; dotted red line indicates tumor boundary (hop = 0); dot size indicates total number of analyzed Visium spots; dot color indicates peak side (blue, stroma; red, tumor). **d,** Peak signal hop analysis across nine matched primary tumor sections, as in c.**e,f,** Boundary-distance decay plot across **e,** 12 metastatic sections and **f,** matched 9 primary sections. Colored lines indicate per-sample smoothed LIMES (top10) UCell score normalized to the boundary mean; gray band indicates ±1 SD. **g,** Masson’s trichrome staining of FFPE lung metastatic sections from ES patients ES2 and ES3, corresponding to patients included in the Visium cohort. Cell nuclei stain black-blue; cytoplasm and muscle fibers, red; collagen and connective tissue, blue-green. Tumor tissue regions are labeled; dashed lines and arrowheads indicate areas of collagen deposition at the tumor-lung interface. Tissue annotation performed by a pathologist in HALO. Scale bar, 100 µm. **h,** As in g, for OS patients OS4, OS5, and OS6.

Our scRNA-seq analysis identified enrichment of a lung fibrosis gene program in AT2 progenitor cells and fibroblast metabolic pathway activity in PATS upon co-culture (Figure 4b), indicating that the LIMES-associated transcriptional response encompasses early fibrotic signaling at the epithelial level. As aberrant epithelial remodeling represents the initiating step of a wound-healing cascade that, when unresolved, proceeds through fibroblast recruitment and collagen deposition toward progressive tissue fibrosis, we asked whether the transcriptional state captured by LIMES is accompanied by histological evidence of fibrotic remodeling at the metastatic interface in patient tissue [17]. To address this, we performed Masson’s trichrome staining - a histochemical method that selectively labels collagen fibers, providing a direct readout of tissue fibrosis - on lung metastatic sections from ES patients ES2 and ES3 and OS patients OS4, OS5, and OS6, corresponding to patients included in the Visium cohort (Supplementary Table 5). Pronounced collagen deposition, visualized as blue-stained connective tissue in proximity to tumor cells, was evident in the peri-tumoral lung parenchyma in both entities (Figures 6f, g), further supporting the notion that infiltrating bone sarcoma cells initiate a fibrotic cascade accompanied by active remodeling of the lung epithelium. Collectively, these findings establish MESCUL as a faithful and informative model of sarcoma-induced lung remodeling: the drastic morphological changes and epithelial remodeling program identified upon bone sarcoma cell contact are transcriptionally recapitulated at the tumor–lung interface in patient metastases, and the collagen deposition observed by Masson’s trichrome staining positions LIMES and its downstream consequences as the histologically manifest onset of a fibrotic cascade in vivo (Figure 6h).

## Discussion

Metastatic homing to the lung has conventionally been viewed as a predominantly tumor-driven process, with the lung epithelium as a passive recipient of infiltrating cells. Our findings reframe this view: the lung epithelium is not a passive substrate but an active participant, and the colonization process is bidirectional. Bone sarcoma cells do not merely invade a receptive tissue, but their arrival triggers rapid, reproducible transcriptional reprogramming toward focal adhesion assembly, ECM remodeling, and a wound-healing state that prepares the parenchyma for tumor infiltration. This response requires direct physical contact; paracrine signaling alone is insufficient to initiate it. The metastatic niche is actively co-constructed at the moment of tumor–epithelial engagement, with implications for when and how therapeutic intervention might interrupt metastatic homing.

The contact-dependent nature of LIMES identifies direct tumor-epithelial engagement as the necessary initiating event. Importantly, this remodeling is not a generic reaction to particulate stimuli or mechanical perturbation: exposure of BME-released lung organoids to polystyrene nanoparticles in the same 3D format activates downstream signaling (pAKT, pERK) but does not induce the morphological spreading or transcriptional reprogramming characteristic of LIMES [30], distinguishing LIMES from a nonspecific stress response and implicating cancer cell-derived molecular signals as the specific trigger, rather than physical contact per se. MESCUL revealed that this program is consistent across both ES and OS and is reproducible across multiple organoid donor backgrounds, suggesting it reflects a stereotyped lung epithelial response to bone sarcoma infiltration rather than a tumor-entity-specific phenomenon. This program encompasses induction of focal adhesion assembly, integrin-mediated ECM remodeling, and emergence of a KRT8⁺ damage-associated transitional state characteristic of aberrant alveolar repair. A wound-healing and fibrotic program at the OS-lung metastatic interface has previously been described primarily in murine models [10]; our spatial transcriptomic and histological profiling of human patient metastases provides the first direct evidence of an equivalent spatially restricted epithelial remodeling response in human tissue, and extends it to ES for the first time. The independent detection of the FN1–integrin axis in human OS tissue by an orthogonal cross-species atlas approach [25] lends further support to the mechanistic model proposed here.

The role of tumor-derived FN1 at the OS–lung interface had been independently anticipated: Reinecke et al. identified FN1 as upregulated in OS lung metastases [10], and Budhathoki et al. nominated it as the dominant metastasis-specific ligand engaging lung epithelial receptors [29]. Our data confirm and mechanistically extend this picture. FN1 is upregulated in co-cultured tumor cells and localizes to the tumor–organoid interface; exogenous FN1 alone is sufficient to trigger organoid remodeling, providing direct functional evidence that FN1 acts as a principal driver of epithelial remodeling. Notably, lung organoid-conditioned medium enriches for factors that induce FN1 expression in sarcoma cells (including GDF15), suggesting the lung microenvironment may prime arriving tumor cells toward this FN1-expressing state. Downstream of integrin engagement, our data identify FAK as the critical mechanosensing transducer, with FAK inhibition via defactinib markedly reducing AREG expression, depleting KRT8⁺ transitional cells, and reducing organoid reshaping. AREG, induced downstream of FAK, acts as an intra-epithelial amplifier; since AREG has been shown to induce FN1 in human lung epithelial cells via EGFR–JNK–AP-1 signaling [31], its induction may sustain epithelial FN1 production independently of continued tumor contact - creating a contact-initiated, self-amplifying feedforward loop that propagates the response across the broader epithelial field. Concomitant upregulation of *MMP1*, *MMP3*, and *MMP10* equips the parenchyma to dissolve ECM barriers. Importantly, this entire signaling architecture operates consistently across ES and OS, extending a mechanism previously inferred only in osteosarcoma to Ewing sarcoma for the first time.

The epithelial origin of the LIMES-associated fibrotic program is consistent with the emerging concept that pathological fibrosis is initiated not by fibroblasts, but by aberrant epithelial–mesenchymal transition [32]. Under conditions of unresolved injury, alveolar epithelial cells arrest in a KRT8⁺ transitional state rather than completing the AT2-to-AT1 differentiation trajectory; this arrest is now recognized as the cellular initiating event of fibrosis, preceding fibroblast recruitment and collagen deposition [14–16]. In the context of bone sarcoma lung metastasis, our data indicate that the FN1–integrin–FAK axis is the contact-dependent trigger of this arrest: FAK inhibition depletes KRT8⁺ PATS and prevents organoid expansion, placing the onset of the fibrotic cascade at the moment of direct tumor–epithelial contact.

The fibrogenic transcriptional program identified by GSEA in lung epithelial cells upon direct sarcoma contact in MESCUL prompted histological evaluation of patient lung metastases, as collagen deposition represents the structural downstream consequence of the epithelial reprogramming observed at the transcriptional level in vitro [17]. Fibrotic stromal changes in bone sarcoma have previously been characterized as a histopathological correlate of chemotherapy-induced tumor necrosis at the primary tumor site [33], and fibrosis has not been regarded as an intrinsic feature of ES progression before [34]. The collagen deposition identified here is spatially confined to the tumor–lung interface, a distribution inconsistent with diffuse therapy-induced stromal remodeling and is transcriptionally predicted by the fibrogenic epithelial reprogramming observed in treatment-naive organoid co-cultures. Notably, EWS-FLI1, the oncogene driving ES pathogenesis, directly suppresses lysyl oxidase (LOX), the enzyme responsible for collagen crosslinking and extracellular matrix stabilization [34], excluding a tumor cell-autonomous origin for the interface-restricted collagen deposition and implicating host lung epithelial remodeling as the principal source. These findings define a contact-dependent mechanism of stromal fibrogenesis at the metastatic niche that is molecularly distinct from therapy-associated stromal reactivity, which is well captured by MESCUL.

Prior spatial transcriptomic analyses of sarcoma have characterized the immune and stromal architecture of primary ES tumors [35] and the OS lung metastatic microenvironment, identifying regionally distinct immunosuppressive niches with fibrocollagenous peritumoral encapsulation [36], aberrant wound-healing programs and KRT8⁺ epithelial intermediates at the metastatic boundary [10], and tumor-derived FN1 as the dominant ligand engaging epithelial integrin and syndecan receptors [25]. We here report a pronounced, spatially restricted enrichment of the LIMES signature at the immediate tumor–lung interface in both OS and ES metastatic lesions; boundary-distance analysis confirmed a stromal-side, boundary-proximal peak in 10 of 12 individual metastatic sections.

Our data provide a first proof of concept that the FAK–AREG axis is pharmacologically targetable; whether this translates into a valid therapeutic strategy for interfering with metastatic colonization requires further investigation. FAK inhibition with defactinib markedly reduced AREG expression, depleted KRT8⁺ transitional cells, and prevented organoid reshaping, providing a first indication that the LIMES program is pharmacologically targetable.

MESCUL is intentionally minimalistic, focusing on the direct interaction between tumor and lung epithelial cells. While this reductionist approach enabled mechanistic exploration of the tumor–epithelium interface, the contribution of immune and stromal compartments, not yet addressed, represents a natural next step toward fully recapitulating the metastatic microenvironment. Spatial transcriptomic findings, while constrained by tissue availability, provide interface-restricted evidence consistent with the organoid observations and a foundation for validation in larger patient cohorts. Beyond bone sarcomas, the consistency of the LIMES program across two genetically distinct tumor entities raises the intriguing possibility that the observed bi-directional tumor-lung epithelium interaction may be of more general relevance to pulmonary metastasis. To our knowledge, an equivalent fibrogenic host response has not been described for lung metastases of other cancer types, suggesting that stromal fibrogenesis at the metastatic niche may represent a sarcoma-specific feature of lung colonization. The extent to which the FAK–AREG axis and the broader LIMES program extend to other tumor entities with a propensity for lung metastasis is an open and clinically relevant question. Beyond the specific findings reported here, MESCUL fills a methodological gap in the field, providing a versatile, human-relevant platform for mechanistic and therapeutic interrogation of bone sarcoma lung metastasis.

## Funding

This research was funded in whole or in part by the Austrian Science Fund (FWF) [grant DOI 10.55776/P35353] and the Federal Ministry of Education, Science and Research (BMBWF). For open access purposes, the author has applied a CC BY public copyright license to any author-accepted manuscript version arising from this submission. Furthermore, the authors would like to acknowledge the following funding sources for their support: Alex’s Lemonade Stand Foundation for Childhood Cancer (20-17258; to M.F., F.H., and H.K.), Cancer Research UK (DRCPFA-Nov24/100001; to F.H.), and St. Anna Kinderkrebsforschung GmbH.

## Author Contributions

Conceptualization: M.M.Z., H.K., B.R.S.; Data curation: T.C.; Formal analysis: M.M.Z., T.C., G.M.B., C.H., N.B., L.W., C.T., B.R.S.; Funding acquisition: F.H., H.K.; Investigation: M.M.Z., N.B., V.S., B.R.S.; Methodology: M.M.Z., U.M., M.v.d.W., B.R.S.; Project administration: H.K., B.R.S.; Resources: F.R., M.W., M.D., K.S., M.v.d.W., M.G.S., B.L.A., M.M., F.H., H.K.; Software: T.C., G.M.B., C.H., C.T.; Supervision: M.F., H.S., A.M., M.M., P.K., F.H., H.K., B.R.S.; Visualization: M.M.Z., T.C., G.M.B.; Writing – original draft: M.M.Z., B.R.S.; Writing – review & editing: M.M.Z., T.C., G.M.B., M.W., C.H., P.K., F.H., H.K., B.R.S.

## Acknowledgements

The authors thank the Core Facility Imaging at the Medical University of Vienna for support with microscopy. Scientific illustrations were commissioned from Ella Maru Studio. Manuscript preparation was assisted by Claude (Anthropic); the authors are responsible for all content.

## Methods

### Ethics statement

The collection of patient tissue for lung organoid generation and tumor cell isolation was conducted in accordance with the guidelines approved by the Ethics Committee of the Medical University of Vienna (EK Nr: 2414/2020).

### Tissue dissociation

Tissue dissociation was performed as previously described with minor modifications [37]. Cryopreserved tissue was thawed at 37°C, transferred to a 15 mL conical tube, and washed by slow addition of 5–10 mL Advanced DMEM/F12 supplemented with 1× Penicillin/Streptomycin, 1× GlutaMAX, and 10 mM HEPES (hereafter AdvDMEM+++) under constant agitation. Fresh tissue was transferred directly to AdvDMEM+++ without prior thawing. Samples were centrifuged at 400 × g for 5 min at 4°C, and pellets were resuspended in digestion medium containing 1 mg/mL collagenase type XI, 10 µM Y-27632, and 50 µg/mL Primocin. Digestion proceeded at 37°C for 1–2 h with periodic inversion, after which the suspension was mechanically dissociated using a 5 mL serological pipette and passed through a 100 µm cell strainer. The filtrate was centrifuged at 400 × g for 5 min at 4°C and washed three times with AdvDMEM+++. Where erythrocyte contamination was evident after the first wash, 2 mL of Red Blood Cell Lysis Buffer (Roche, cat. no. 11814389001) was applied for 5 min at room temperature, followed by three additional washes. All cell cultures were routinely tested for mycoplasma contamination using the MycoAlert Mycoplasma Detection Kit (Lonza).

### Lung organoid culture

Patient-derived lung organoids were established according to Sachs et al.[13]. Briefly, following tissue dissociation, the cell pellet was resuspended in ice-cold basement membrane extract (BME; Cultrex growth factor-reduced BME type 2, Trevigen, 3533-010-02) at a ratio of 70% BME to 30% residual AdvDMEM+++. The suspension was plated as 10–15 µL domes per well in pre-warmed 24-well plates, inverted, and solidified at 37°C for 20 min. Lung organoid medium (LO medium, according to Sachs et al.) supplemented with 5 µM Y-27632 was added; Y-27632 was omitted after the first three days. Medium was refreshed every 3–4 days. After 14–21 days, organoids were passaged at a 1:2–1:6 ratio by mechanical dissociation through a 27G needle (5–10 passes in 1 mL ice-cold AdvDMEM+++) and filtered through a 40 µm strainer to retain a single-cell suspension. Cells were centrifuged at 400 × g for 5 min at 4°C, resuspended in cold BME, and re-plated as described above.

### MESCUL co-culture setup

To initiate co-cultures, lung organoids were recovered from BME 7-14 days after passage by resuspending domes in ice-cold AdvDMEM+++ followed by centrifugation. Organoid concentration was determined by brightfield microscopy.

Tumor cells were obtained either directly from tissue dissociation (as described above) or by mechanical disruption of patient-derived tumoroids, as previously described [38,39]. Following dissociation to single-cell suspension, cell viability and concentration were determined by Trypan Blue exclusion using a Countess III automated cell counter (Thermo Fisher Scientific). Cells were subsequently labeled with CellTrace™ Violet or CellTrace™ CFSE (Invitrogen, cat. no. C34557/C34554) according to the manufacturer’s instructions prior to co-culture.

Organoids and tumor cells were plated on tissue culture plates without exogenous extracellular matrix support. Co-cultures were maintained in a defined medium based on the lung organoid medium described by Sachs et al.[13], supplemented with 1× B27, 1× Penicillin/Streptomycin, and 1× GlutaMAX.

For continuous live-cell imaging, co-cultures were monitored using an Olympus IX83 widefield microscope (Core Facility for Imaging, Medical University of Vienna) or an Incucyte S3 Live-Cell Analysis System (Sartorius). For defined-timepoint acquisition of brightfield and fluorescence images, co-cultures were imaged on a Leica TCS SP8X confocal microscope. Images were acquired using LAS X software (Leica Microsystems).

For barrier co-culture experiments, organoids were plated in the upper chamber of Incucyte Clearview 96-well Chemotaxis Plates (Sartorius, cat. no. 4582; paired with Reservoir Microplate, cat. no. 4601) in co-culture medium. Tumor cells were added either to the upper chamber alongside organoids (direct co-culture) or to the lower reservoir chamber only (indirect co-culture). Monoculture controls received co-culture medium in the lower chamber without tumor cells.

Organoid area was quantified from brightfield images using Fiji/ImageJ [40] by manual region-of-interest delineation per organoid. Area values (µm²) were log₁₀-transformed prior to statistical modelling to account for right-skewed size distributions. For longitudinal within-experiment comparisons in which the same individual organoids were tracked across two timepoints (Figure 2d, e), statistical comparisons were performed using the Wilcoxon matched-pairs signed-rank test (two-tailed), treating each organoid as its own matched pair. For multi-condition comparisons involving a nested data structure, linear mixed-effects models were fitted using the lme4 and lmerTest packages in R, with the random-effects structure adapted to the experimental design. For the three-condition transwell experiment comprising three independent biological replicates (independent co-culture experiments initiated on separate days) and two technical replicate wells per condition per replicate (Figure 2i), the model was specified as: log₁₀(area) ∼ condition + (1 | experiment/well). For individual single-replicate co-culture experiments each comprising four technical replicate wells per condition (Figure 3a-d), the model was reduced to: log₁₀(area) ∼ condition + (1 | well), as no between-experiment variance was present. For the FN1 treatment experiment, organoids from two independent patient-derived lines (L-EWS-050, L-EWS-070) were assessed across two independent experimental replicates per line, each seeded from independent organoid passages (replicate 1: 4 wells per condition; replicate 2: 3 wells per condition). To confirm the FN1 effect within each donor line independently (Supplementary Figure 2c, d), a per-line model was fitted as: log₁₀(area) ∼ condition + (1 | experiment/well), with Kenward-Roger degrees of freedom. To obtain a combined cross-patient estimate accounting for inter-donor variability (Figure 5f), a joint model was fitted as: log₁₀(area) ∼ condition + (1 | patient) + (1 | batch/well), where batch denotes the patient-by-experiment combination. In all mixed-effects models, pairwise contrasts were computed using the emmeans package with Tukey correction. Effect sizes are reported as estimated fold-changes in median organoid area, back-transformed from the log₁₀ scale, relative to mono-culture or untreated controls. All p-values are two-sided; p < 0.05 was considered statistically significant. Graphs were generated using ggplot2 in R (v4.5.1) and GraphPad Prism (v11.0.0(84)).

### Drug treatment

Lung organoids were recovered from BME and re-plated on 96-well plates (Corning), or co-cultures were established as described above. Compounds were prepared at 2× the intended final concentration and added in equal volume to the culture medium. Cultures were monitored by live-cell imaging (Incucyte) and evaluated by organoid area quantification or immunofluorescence staining as described below. Defactinib dose–response was assessed using CellTiter-Glo® (CTG; Promega, Cat. #G7570) for tumor cells and CellTiter-Glo® 3D (CTG 3D; Promega, Cat. #G9681) for organoids. ES-ZH-002 and ES-ZH-009 cells were seeded in 96-well viewplates (Revvity, Waltham, MA, USA, Cat. #6005181) one day prior to treatment. Lung organoids were recovered from BME 11 days after passage by resuspending domes in ice-cold AdvDMEM+++ followed by centrifugation (400×g, 5 min, 4°C), and plated into 96-well viewplates one day prior to treatment. Defactinib was applied across a ten-point serial dilution spanning four orders of magnitude (tumor cells: 0.051–20,000 nM; organoids: 0.026–10,000 nM), in triplicate (final volume 100 µL per well). After 72 hours at 37°C / 5% CO_2_, 100 µL of CTG reagent (1:4 in DPBS; 10 min shaking, 15 min dark) or undiluted CTG 3D (5–10 min shaking, 30 min dark) was added per well. Luminescence was recorded on a Spark Cyto plate reader (Tecan Group, Männedorf, Switzerland). The final DMSO concentration was ≤0.2%. Raw luminescence was normalized to vehicle controls to express percentage viability, and dose–response curves were fitted using SynFit (synfit.app).

### Fixation and immunofluorescence staining

Cultures were washed once with PBS and fixed in 4% paraformaldehyde (PFA) in PBS for 30 min at room temperature, followed by two PBS washes. Permeabilization was performed in 0.1% Triton X-100 in PBS for 20 min at room temperature, followed by blocking in 1% BSA in PBS with 0.1% Tween-20 for 45 min at room temperature. Samples were incubated with primary antibody in blocking buffer overnight at 4°C, washed three times with PBS, and incubated with the appropriate fluorophore-conjugated secondary antibody in blocking buffer overnight at 4°C. On the third day, samples were washed twice with PBS and incubated with CellMask™ Actin Tracking Stain (Invitrogen, A57249) for 30 min at room temperature. Following two additional PBS washes, samples were mounted in ibidi Mounting Medium (ibidi, 50001). Confocal imaging was performed on a Leica TCS SP8X confocal microscope. Images were acquired using LAS X software (Leica Microsystems) and linearly adjusted for brightness and contrast for display purposes; identical settings were applied to all compared panels. Primary and secondary antibodies are listed in Supplementary Table 6.

### Single-cell RNA sequencing

For scRNA-seq, short-term co-cultures were harvested by passing the suspension through a 40 µm strainer to separate organoid-associated tumor cells (retained on the strainer) from unassociated tumor cells (flow-through). Loosely adherent cells that had settled on the well bottom were detached by vigorous pipetting and pooled with the organoid-associated fraction. Both fractions were centrifuged at 400 × g for 5 min at 4°C and resuspended in Accutase. The flow-through fraction was dissociated for 3–5 min; the organoid-associated fraction required 15–25 min. Cell viability and counts were assessed by Trypan Blue exclusion using a Countess III automated cell counter (Thermo Fisher Scientific). Single-cell suspensions were processed for scRNA-seq using the 10x Genomics Chromium Single Cell 3′ Gene Expression Kit v3.1 according to the manufacturer’s instructions. Monocultures of lung organoids and tumor cells were processed in parallel. Library quantity and quality were assessed using the Qubit Fluorometer with the High Sensitivity dsDNA Assay Kit (Thermo Fisher Scientific) and the Agilent TapeStation with High Sensitivity D5000 ScreenTape (Agilent Technologies). Libraries were sequenced by Novogene (partial lane, NovaSeq X Plus, PE150), targeting 10 000 cells per sample.

### Raw data processing, quality control, and filtering

Raw sequencing data were processed with the CellRanger multi v9.0.1 software (10x Genomics) for cell-level demultiplexing and alignment to the human reference transcriptome (refdata-gex-GRCh38-2024-A assembly provided by 10x Genomics). We used the filtered featured count matrices for all downstream analysis steps. All preprocessing and analyses steps except differential expression analysis were performed in Python (3.12.3) using the scanpy [41] (1.11.1) and additional packages [anndata (0.11.4), decoupler-py (2.0.2), gseapy (1.1.8), matplotlib-base (3.10.1), numpy (2.2.6), pandas (2.2.3), scikit-learn (1.5.2), scipy (1.13.1), seaborn (0.13.2), statsmodels (0.14.4)]. In order to remove cells that may be stressed or dying, and thus deviate significantly from the norm or have low gene/UMI counts (empty droplets), we performed the following filtering steps on the UMI count matrix per batch: 1) remove cells with mitochondrial count fraction > 25%, 2) remove cells with less than 800 genes detected, 3) remove cells with z-scored log10-transformed transcript number outside the range [-3, 3], 4) remove cells with unusually low/high gene count relative to transcript count (loess fit to the relationship of the log10-transformed values of number of genes as function of transcripts; cells with z-scored residuals outside the range [-5, 5] removed). To integrate our five batches we merged the data, normalized, log[FH1.1]-transformed and performed PCA with 50 principal components on 5000 highly variable genes. Based on the PCA we used the batch-balanced K-Nearest-Neighbour (bbKNN) algorithm [42] from scanpy to obtain a KNN-graph that was adjusted for batch effects and used for subsequent Leiden clustering [43] with resolution=0.7 and UMAP[44] representation.

### Cell type annotation and label transfer

To annotate the identified clusters with cell type labels, we used scPoli [45] from the scarches package (0.6.1) to map our data and the Human Lung Cell Atlas [19] (only healthy samples[FH2.1]) into a common space and transfer the labels of the atlas to our data using the inbuilt prediction function of scPoli. We manually refined the resulting cell type annotations by using a set of curated marker genes (Basal cells: *TP63*, *KRT5*; Activated basal: *PDPN*; Secretory: *SCGB1A1*, *CYP2F1*, *MUC5B*; Alveolar: *KRT8*, *KRT18*, *EPCAM*; Alveolar type 1 (AT1): *CAV1*; Alveolar type 2 (AT2): *CLIC5*, *HOPX*, *SFTB*, *SLC34A2*; Ciliated: *TUBA1A*, *RSPH1*; Proliferating: *MKI67*).

### Differential expression analysis

To identify differentially expressed genes between co-cultured and monocultured lung organoids, we used DESeq2 [46] (1.44.0) in R (4.4.1). We first generated pseudobulks per cell type and batch by summing the raw counts and then fit DESeq2 models on the pseudobulks, while controlling for batch effects using the following design: ∼ condition + batch, where condition refers to the culture condition. DESeq2 was used with the Wald test and default parameters[FH3.1].[FH4.1] Genes with an adjusted p-value < 0.05 and a log2 fold change > 1 were defined as significantly upregulated in the co-culture. P-values were adjusted for multiple testing using the Benjamini-Hochberg procedure [47].

### Gene set enrichment analysis

To test for enriched gene sets in response to co-culture with tumor cells, we used pre-ranked gene set enrichment analysis (GSEA) with the results from the pseudobulk differential expression test compared to the monoculture. We used the Wald statistic (cf. differential expression analysis above) to rank the genes. Gene set enrichment tests were performed per cell type using the prerank function of the GSEApy [48] package (1.1.8) with 1000 permutations. Gene sets were taken from Gene Ontology - Biological Process [49,50] 2023, KEGG [51] 2021, Reactome [52] 2022 and WikiPathways [53] 2023 obtained via Enrichr [54].

### Definition of the Lung Interface Metastasis Signature (LIMES)

To define LIMES, a curated gene signature describing the molecular response of the lung organoids to co-culture with bone sarcoma cells, we again performed differential expression analysis with the cell types (AT2, AT2 progenitors, proliferating alveolar cells, activated basal cells and PATS) that showed enrichment of gene sets of ECM remodeling, focal adhesion, and cytoskeletal programs. The differential expression analysis was performed as described above but with the following design: ∼ condition + batch + celltype, to obtain a cell type and batch independent effect for the co-culture condition. The resulting list of differentially expressed genes (adjusted p-value < 0.05) was filtered (baseMean>= 10 to remove spurious genes with low expression) and ranked by log2 fold change. The top 10 genes with the highest log fold change are the LIMES signature (*FSCN1, MMP1, KRT6A, AREG, EREG, CD274, SLC7A5, BCAT1, CCL5, DSG3*) (Supplementary Table 3).

### Cell-cell communication analysis

To identify the receptors and downstream targets of FN1 we leveraged the NicheNet [55] database[FH5.1] (accessed via [56]). We used the regulatory potential score of NicheNet to identify top-ranking receptors and downstream that were detected in our scRNA-seq dataset.

### Spatial transcriptomics**-** Library preparation and sequencing

Formalin-fixed, paraffin-embedded (FFPE) tissue sections (5 µm) from matched primary tumors and lung metastases of three ES and six OS patients (Supplementary Table 5) were processed using the 10x Genomics Visium Spatial Gene Expression workflow for FFPE (v1 chemistry) according to the manufacturer’s protocol. Hematoxylin and eosin (H&E)-stained sections were imaged at the Core Facility for Imaging at the Medical University of Vienna prior to library preparation and used for pathologist-guided tissue annotation. Visium libraries were sequenced at the Genomics Facility, Medical University of Vienna on Illumina NextSeq500 and NextSeq2000 instruments, targeting a minimum of 25,000 reads per spot.

### Spatial transcriptomic **-** Data processing

Spatial transcriptomics data were processed with Space Ranger (v2.0.1, 10X Genomics) against the standard GRCh38 reference genome with the count function for each sample. Individual samples were joined in a single Seurat object and counts were log normalized using Seurat’s NormalizeData function with default parameters. LIMES signature score was calculated on normalized counts using the function AddModuleScore_UCell from the UCell package [57] (v2.8.0). Pathologist annotations were exported from QuPath [58] (v0.6) as GeoJSON files and transferred to Seurat objects by spatial join, using spot centroid coordinates in full-resolution pixel space as the reference frame. Annotation conflicts between overlapping regions were resolved by prioritising the annotation with the smallest polygon area, favouring histologically specific labels over broad tissue categories.

To quantify gene expression gradients relative to the tumor boundary, we defined boundary spots as tumor-annotated spots with at least one non-tumor neighbour within 1.4× the estimated spot pitch, as determined empirically from the median nearest-neighbour distance in physical coordinate space. Hop distance was then computed using multi-source breadth-first search (BFS) initialised simultaneously from all boundary spots, assigning each spot the minimum number of spot hops to the nearest boundary spot. Spots within the tumor received negative hop values and spots in the surrounding tissue received positive values, with boundary spots assigned zero. This approach leverages the regular hexagonal geometry of the Visium array and is independent of physical coordinate scaling, making it directly comparable across experiments regardless of scanner resolution.

For cohort-level analysis, inter-sample variation in baseline expression was removed by subtracting the mean signal at boundary spots (hop 0) from each sample independently, yielding a boundary-normalised expression value. Per-sample decay profiles were computed by binning spots by hop distance and calculating mean, median, standard deviation, and standard error per bin. The peak hop distance was defined as the hop bin with maximal smoothed normalised expression across all bins. All analyses were performed in R using the Seurat [59] (v5.3.1), sf [60](v1.0-17), and ggplot2 [61] (v3.4.4) packages.

### Masson’s trichrome staining

For the Masson’s trichrome staining, the MASSON’s Trichrome with Aniline Blue staining kit (Morphisto, 18156.00500) was used on 5 µm FFPE tissue sections. The protocol was performed according to the supplier’s instructions.

In the final steps, slides were mounted with Entellan (Sigma-Aldrich, 1.07960) in a fume hood, dried overnight, and acquired in brightfield mode using a Vectra Polaris 1.0 scanner at 20× magnification. Tissue annotations were provided by an expert pathologist and were used accordingly for analysis.

### Cytokine Array

The secretome of lung organoid conditioned medium was profiled using the Proteome Profiler Human XL Cytokine Array Kit (R&D Systems, ARY022B). Lung organoid conditioned medium was collected after 72 hours incubation, centrifuged at 300 × *g* for 5 min to remove debris, and used undiluted. Array membranes were processed according to the manufacturer’s protocol. Briefly, membranes were blocked, incubated overnight at 4 °C with sample, washed, and incubated with a biotinylated detection antibody cocktail followed by streptavidin–HRP. Chemiluminescent signal was detected using a Bio-Rad ChemiDoc imaging system. Spot pixel densities were quantified using the Protein Array Analyzer macro for Fiji (G. Carpentier, Université Paris Est Créteil; available at http://image.bio.methods.free.fr/ImageJ/)[40]. Raw pixel densities were normalized to the mean of the array reference spots on each membrane. Background was determined by measuring parallel membranes incubated with organoid medium alone and subtracted spot-by-spot from conditioned medium values. Factors with a background-subtracted net signal ≥ 0.1 (relative to reference spots) were considered detected.

## Data and code availability

Single-cell RNA-seq data generated in this study will be deposited in the European Genome-Phenome Archive (EGA) prior to publication. Computer code used for data analysis will be shared via our GitHub organization (https://github.com/cancerbits).

## Notes

### Competing Interest Statement

The authors have declared no competing interest.

