## Supplementary figures and images for "A human lung organoid co-culture model of early bone sarcoma metastasis reveals contact-dependent epithelial remodeling at the metastatic interface"

### Supplementary Movie 2

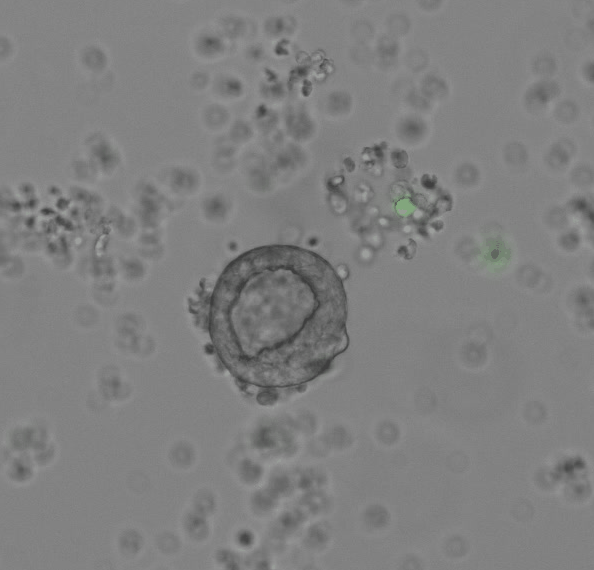
